# Inhibition of protein or glutamine biosynthesis affect the activation of the light stimulated SBiP1 chaperone by dephosphorylation in Symbiodiniaceae

**DOI:** 10.1101/2024.08.01.606268

**Authors:** Raúl Eduardo Castillo-Medina, Tania Islas-Flores, Estefanía Morales-Ruiz, Marco A. Villanueva

**Affiliations:** Instituto de Ciencias del Mar y Limnología, Unidad Académica de Sistemas Arrecifales, Universidad Nacional Autónoma de México-UNAM, Prol. Avenida Niños Héroes S/N, Puerto Morelos, Quintana Roo 77580, México

**Keywords:** chaperone, light stimulation, phosphorylation, phototransduction, protein synthesis, Symbiodiniaceae

## Abstract

Phosphorylation/dephosphorylation is fundamental for transduction of external stimuli into physiological responses. In photosynthetic dinoflagellates *Symbiodinium microadriaticum* CassKB8, Thr-phosphorylated SBiP1 under dark conditions, is activated through dephosphorylation upon light stimuli. We evaluated the effect of protein synthesis inhibitors on light modulated Thr phosphorylation of SBiP1. Inhibition of cytoplasmic protein synthesis by cycloheximide but not of chloroplastic protein synthesis by chloramphenicol, promoted inactivation via Thr re-phosphorylation of the protein under the light. Additionally, inhibition of glutamine synthetase by glufosinate produced a delay in the light induced activation by dephosphorylation of the chaperone. Heat shock reverted the effect in cycloheximide-treated cells suggesting that heat stress overrides the cycloheximide-induced inactivation to restore chaperone activity. These results suggest that light and stress are critical switches of SBiP1 chaperone activity that function along with common pathways of protein synthesis and ammonia assimilation, and further confirm that the light induced SBiP1 Thr dephosphorylation is independent of photosynthesis.

## Introduction

The marine photosynthetic dinoflagellate *Symbiodinium microadriaticum* CassKB8 (henceforth CassKB8) was originally isolated from the jellyfish *Cassiopea xamachana* in its endosymbiotic stage and has been widely used as a model of study in its *in vitro* cultured stage [1,2,3,4–6]. These microorganisms can either live freely in the water column or as endosymbionts of a variety of cnidarian hosts. Since they respond to diurnal/nocturnal photoperiod cycles, they require the necessary signal-transduction machinery to adapt their responses depending on the environmental conditions.

Protein kinases and phosphatases are responsible for the exquisite regulation of key target molecules through phosphorylation and dephosphorylation within cellular signal-transduction cascades and a significant representation of kinase sequences has been reported in Symbiodininaceae [2,7]. Among such target molecules reported to be modulated by phosphorylation, are the BiP chaperones [5,6,8–10] resident of the Endoplasmic Reticulum (ER). In this compartment, they assist in the folding and assembly of newly synthesized proteins as they undergo translocation into the ER, and they can also associate with misfolded and/or underglycosylated proteins [8,11].

Post-transcriptional phosphorylation of BiP results in profound intracellular functional changes; for example, phosphorylated BiP is believed to assemble into oligomers as the inactive form whereas only the monomeric active dephosphorylated form can bind to clients [8,12]. Thus, phosphorylation/dephosphorylation of BiP correlates with inactivation/activation, respectively, of the chaperone activity [5,6,8,10,12]. This inactivation was observed as an increase in CrBiP phosphorylation in the presence of the inhibitor of protein synthesis cycloheximide (CHX) in *Chlamydomonas reinhardtii* [10]. The conclusion was that the CrBiP phosphorylation state was indirectly regulated by TOR (Target of Rapamycin) through the control of protein synthesis after a screen for targets of TOR in this alga [10].

Recently, we reported the light modulated phosphorylation/dephosphorylation of a BiP-like protein, SBiP1 from CassKB8 [5,6]. The presence of SBiP1 sequences was observed in all Symbiodiniaceae species reported so far [6], but we have focused on determining its properties from cultured CassKB8 which naturally occurs as the endosymbiont of *C. xamachana*. SBiP1 was shown to be highly phosphorylated on Thr during the 12 h of the CassKB8 dark cycle and but upon shifting to light, Thr phosphorylation decreased within 30 min [6]. Dephosphorylation occurred independent of photosynthesis and the wavelength of the visible light spectra but was highly sensitive to light intensity as it occurred with as low as a 1 µmole photon m^-2^ s^-1^ stimulus [6]. Furthermore, in highly phosphorylated SBiP1 in the absence of light, the stressful temperature of 32°C for this species also induced its Thr dephosphorylation [5]. These data suggested activation/inactivation of the chaperone function of SBiP1 by regulation of its Thr phosphorylation levels under homeostasis and/or stress conditions in CassKB8.

To gain further knowledge on the relationship of the SBiP1 modulation of by phosphorylation with cellular events, we evaluated the effect of protein synthesis and glutamine synthetase inhibition on the phosphorylation status of the protein and thus confirm our hypothesis that a correlation of activation/inactivation of the chaperone with dephosphorylation/phosphorylation, respectively, exists. We found that activation of SBiP1 was inhibited upon treatment with CHX but not with chloramphenicol (CPL); however, this phenomenon was only delayed by the glutamine synthetase inhibitor, glufosinate (GLUF). Finally, short-term heat shock re-activated the chaperone by reverting the cycloheximide induced inactivation. These results suggest a strong interrelationship between SBiP1 chaperone activation/inactivation and cellular processes in a concerted regulation by nutritional and stress cues in addition to light.

## Results

### SBiP1 phosphorylation on Thr in CassKB8 cells occurs under continuous light in the presence of cycloheximide but not chloramphenicol

Since it has been previously documented that BiP chaperones from *C*. *reinhardtii* are activated by dephosphorylation at the onset of *de novo* protein synthesis, which occurs downstream of the TOR pathway, and the effect is reversed by the protein synthesis inhibitor CHX [10], we tested the effect of this inhibitor on the phosphorylation behavior of SBiP1 under maximum dephosphorylation conditions, starting 3 h after the onset of the light cycle. We observed that the Thr phosphorylation level of SBiP1 (Fig. 1, SBiP1-p, upper panels) was indeed minimum 3 h after the onset of light in the control treated for 30 additional min under light with vehicle alone (Fig. 1A, lane EtOH, upper panel, 30 min; Fig. 1B, white bar, 30 min), and remained unchanged for the next 1, 4 and 7 h after the addition of vehicle (Fig. 1A, lanes EtOH, upper panel, 1, 4 and 7 h, respectively; Fig. 1B, white bars, 1, 4 and 7 h, respectively). In contrast, after 3 h of light and treatment with CHX for further 30 min under light, a clear increase in the Thr phosphorylation level was observed (Fig. 1A, lane CHX, upper panel, 30 min; Fig. 1B, grey bar, 30 min). The increase in phosphorylation was sustained for the next 1 and 4 h after CHX was added (Fig. 1A, lanes CHX, upper panel, 1 and 4 h, respectively; Fig. 1B, grey bars, 1 and 4 h, respectively). This increase in Thr phosphorylation was statistically significant after quantitation by densitometry of the bands and normalization against the total SBiP1 protein detected by anti-SBiP1 antibodies (Fig. 1A, lower panels, SBiP1) from three biological replicates (Fig. 1B, asterisks on grey bars). At the same time, the SBiP1 detection was used as an internal loading control for each respective lane (Fig. 1A, lower panels, SBiP1). After 7 h of treatment with CHX to the cells, the SBiP1 Thr phosphorylation level decreased to statistically non-significant values (Fig. 1B, grey bar, 7 h), although the SBiP1-p band corresponding to phosphorylated SBiP1 (Fig. 1A, lane CHX, upper panel, 7 h) was still observed more intense than in the control (Fig. 1A, lane EtOH, upper panel, 7 h). In parallel, we tested the effect of the chloroplast-specific protein synthesis inhibitor CPL on the level of SBiP1 phosphorylation. We observed no significant effect on the Thr phosphorylation levels of SBiP1 when CPL was added 3 h after the onset of light and the cells further incubated for 30 min, 1, 4 and 7 h (Fig. 1A, lanes CPL, upper panel, 30 min, 1, 4 and 7h, respectively; Fig. 1B, dotted bars, 30 min, 1, 4 and 7 h, respectively). As a positive control for the CPL inhibitory effect on chloroplast protein synthesis, we followed the presence of the photosystem II, D1 protein, throughout the treatment. We observed that, indeed, the protein level diminished as a function of the time of CPL treatment compared to the control (Supp. Fig. 1). These results indicated that global protein synthesis inhibition exerts an effect on light induced SBiP1 dephosphorylation whereas specific chloroplast protein synthesis inhibition does not.

**Figure 1.**
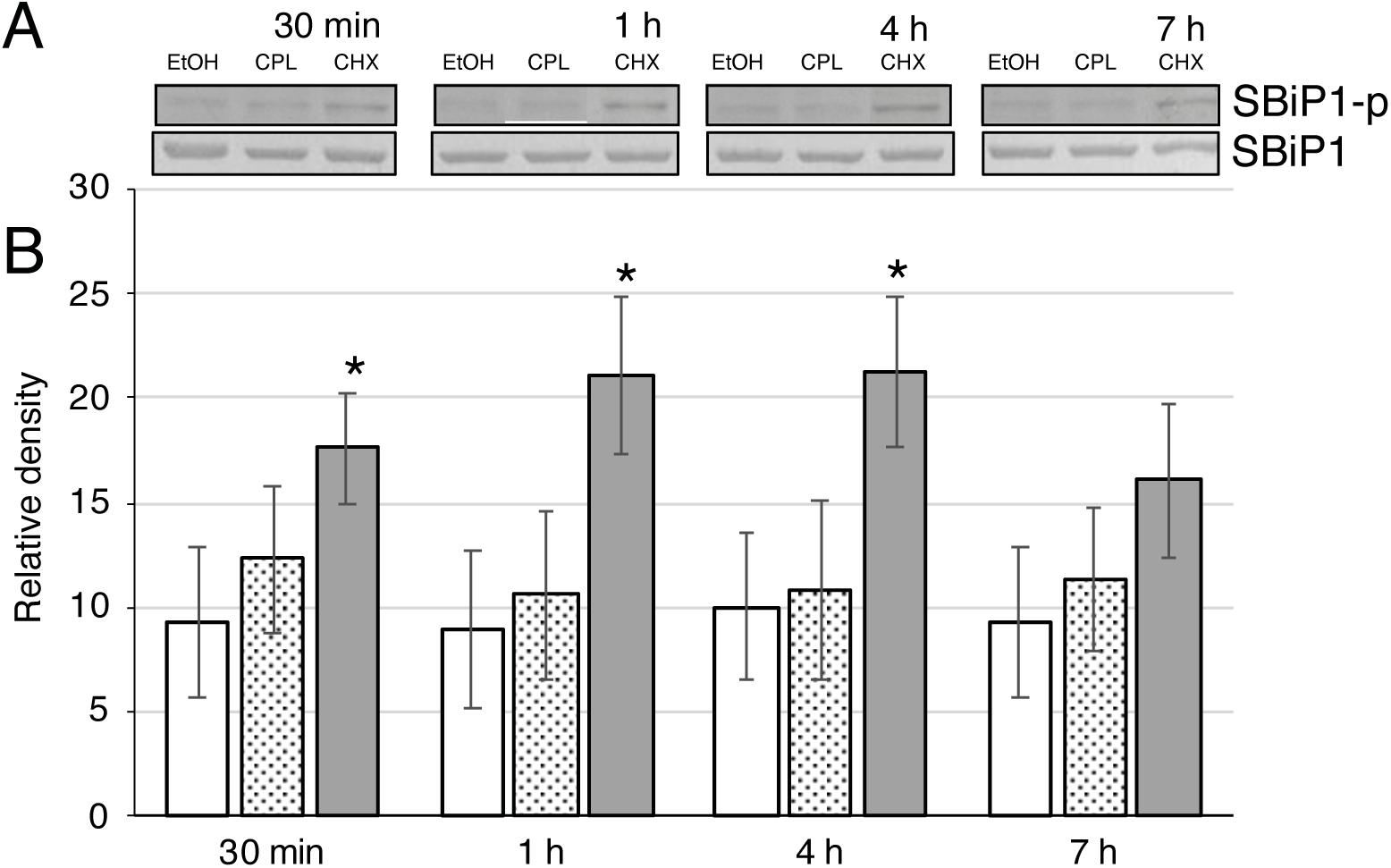
The Thr phosphorylation response of SBiP1 in the presence of light and protein synthesis inhibitors. A. Upper panel: representative western blots showing the level of Thr phosphorylation of SBiP1 (SBiP1-p) from dark-adapted CassKB8 cells in the presence of 0.1 mM cycloheximide (CHX) or 0.1 mM chloramphenicol (CPL) compared with vehicle (EtOH) after light stimulation for 30 min, 1, 4 and 7 h; lower panel: western blot of the same time points analyzed with anti-SBiP1 antibodies (SBiP1) as internal protein load and normalization control for densitometric analysis. B. Densitometric analysis of three biological replicates (± standar error of the mean) from each time point quantitatively showing the levels of SBiP1 Thr phosphorylation after treatments with CHX (grey bars) or CPL (dotted bars), compared with vehicle (white bars). Asterisks show statistically significant values compared to the control for each time (p<0.05).

### High levels of SBiP1 phosphorylation on Thr in CassKB8 cells are sustained during the dark phase and after the transition to light in the presence of cycloheximide

We wanted to assess whether CHX reverted or rather prevented the dephosphorylation of SBiP1 after the transition from darkness to the light phase of growth. Thus, we added CHX under darkness 2 h prior to the onset of the light phase of the growth cycle and monitored the SBiP1 Thr phosphorylation after the switch to light for 30 and 60 min. As previously observed, the high level of Thr phosphorylation of SBiP1 typical of cells under darkness was present in cells treated with vehicle for 2 h prior to the onset of light (Fig. 2A, left upper panel (SBiP1-p), Dark, 12 h; Fig. 2B, black bar, Control). A similar level of Thr phosphorylation was observed in cells treated with CHX for 2 h prior to the onset of light (Fig. 2A, right upper panel (SBiP1-p), Dark, 12 h; Fig. 2B, black bar, CHX). As expected, in vehicle treated cells, the typical dephosphorylation of SBiP1 was observed 30 and 60 min after the onset of light (Fig. 2A, left upper panel (SBiP1-p), lanes 30 and 60 min, respectively; Fig. 2B, left white bars, Control, respectively). Conversely, the SBiP1 Thr phosphorylation level remained with little change in CHX treated cells after light exposure for 30 or 60 min (Fig. 2A, right upper panel (SBiP1-p), lanes 30 and 60 min, respectively; Fig. 2B, light grey bars, CHX, respectively). These results indicated that the presence of CHX in CassKB8 cells under darkness is sufficient to prevent the light induced Thr dephosphorylation of SBiP1 for at least the time tested of 1 h after the onset of light and suggest that light triggers processes that impinge on protein biosynthesis.

**Figure 2.**
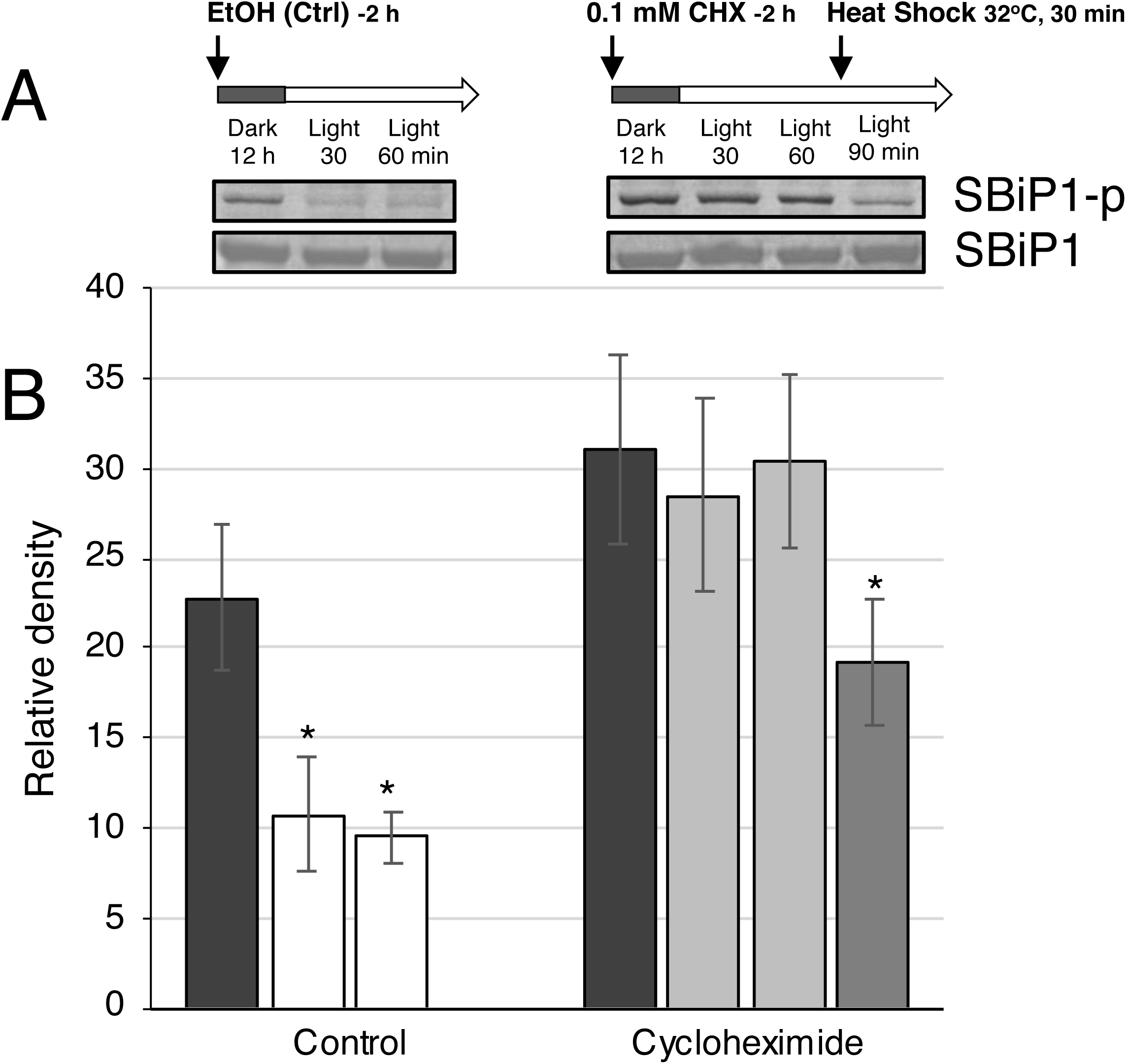
Effect of cycloheximide treatment with or without a heat shock on CassKB8 cells during the dark-to-light transition. A. Upper panels: representative western blots showing the level of Thr phosphorylation of SBiP1 (SBiP1-p) from dark-adapted CassKB8 cells incubated for 2 h before the light stimulation with 10 µM cycloheximide (CHX), or vehicle (EtOH) only, and after 30 and 60 min of light. SBiP1 is highly Thr-phosphorylated after 12 h of continuous darkness regardless of the treatments (SBiP1-p; Dark, 12 h). The SBiP1 Thr phosphorylation levels from dark adapted cells decrease significantly after a 30- or 60-min light stimulus in the presence of vehicle (SBiP-p, left panel; lanes 30, 60 min, respectively); conversely, the SBiP1 Thr high phosphorylation level is sustained at the same time points after light stimulation when CHX is present (SBiP1-p, right panel; lanes 30, 60 min, respectively). SBiP1 Thr high phosphorylation levels (SBiP1-p) sustained after 60 min of light stimulation in the presence of CHX decrease significantly under light after a 30-min heat shock and a 90-min total light stimulus (SBiP1-p, right panel; lane 90 min). Lower panels: western blots of the same treatments and time points analyzed with anti-SBiP1 antibodies (SBiP1) as internal protein loading controls and used for normalization of the densitometric analysis, showing constant levels of the SBiP1 protein throughout the treatments. B. Densitometric analysis of four biological replicates (± standar error of the mean) for each treatment quantitatively showing the levels of SBiP1 Thr phosphorylation under darkness (black bars), or after 30- and 60-min light stimulation in the presence of vehicle (white bars), CHX (light grey bars) or after a 30 min heat shock at 32°C (grey bar). Asterisks show statistically significant values compared to the dark condition for each treatment (p<0.05).

### Short-term heat shock reverts the effect of CHX on Thr dephosphorylation of SBiP1 in CassKB8 cells

One set of the CHX treated cells after 60 min of light stimulation in the presence of CHX were subjected to treatment with 32°C for 30 additional min and then analyzed by western blot with anti-pThr antibodies. We observed that even though CHX prevented the Thr dephosphorylation of SBiP1 after the light stimulus, the heat shock overrode this effect and promoted the dephosphorylation of the chaperone (Fig. 2A, right upper panel (SBiP1-p), 90 min; Fig. 2B, dark grey bar, CHX). The results were statistically significant from three biological replicates (asterisks on bars). This indicated that a short heat shock treatment was sufficient to reactivate the SBiP1 chaperone.

### Light stimulated SBiP1 dephosphorylation on Thr in CassKB8 cells is delayed by the glutamine synthetase inhibitor glufosinate

Since the inhibition of protein biosynthesis appeared to prevent dephosphorylation (thus maintaining the chaperone active state) of SBiP1, we explored the effect of inhibition of another biosynthethic process. We tested the competitive inhibition of glutamine synthetase (which aminates glutamate to produce glutamine) by 1 mg/ml GLUF, on the light stimulated SBiP1 dephosphorylation. GLUF was added to the cell cultures at the onset of the dark cycle and incubated for 12 h in the dark to first determine whether this treatment would affect the viability of the cells at the times assayed. When we checked viability of the GLUF trated cultures by Evans blue [13], we observed that after 12 h in the presence of GLUF, the cells did not show any blue staining (Fig. 3A, middle panel, GLUF) and quantitation showed 100 % viability (Fig. 3C, diagonally hatched bar), similar to the untreated cells (Control in both Fig. 3A, upper panel, and Fig. 3C, white bar, respectively). This indicated that the cells were undisturbed by the treatment. On the other hand, heat-killed cells were both stained blue (Fig. 3A, lower panel, boiled) and less than 50 % viable (Fig. 3C, dotted bar). Cell concentration was ∼ 30 % lower in the treated cells (Fig. 3B, diagonally hatched bar), compared to the untreated cells (Fig. 3B, white bar) indicating a possible delay in cell division induced by GLUF.

**Figure 3.**
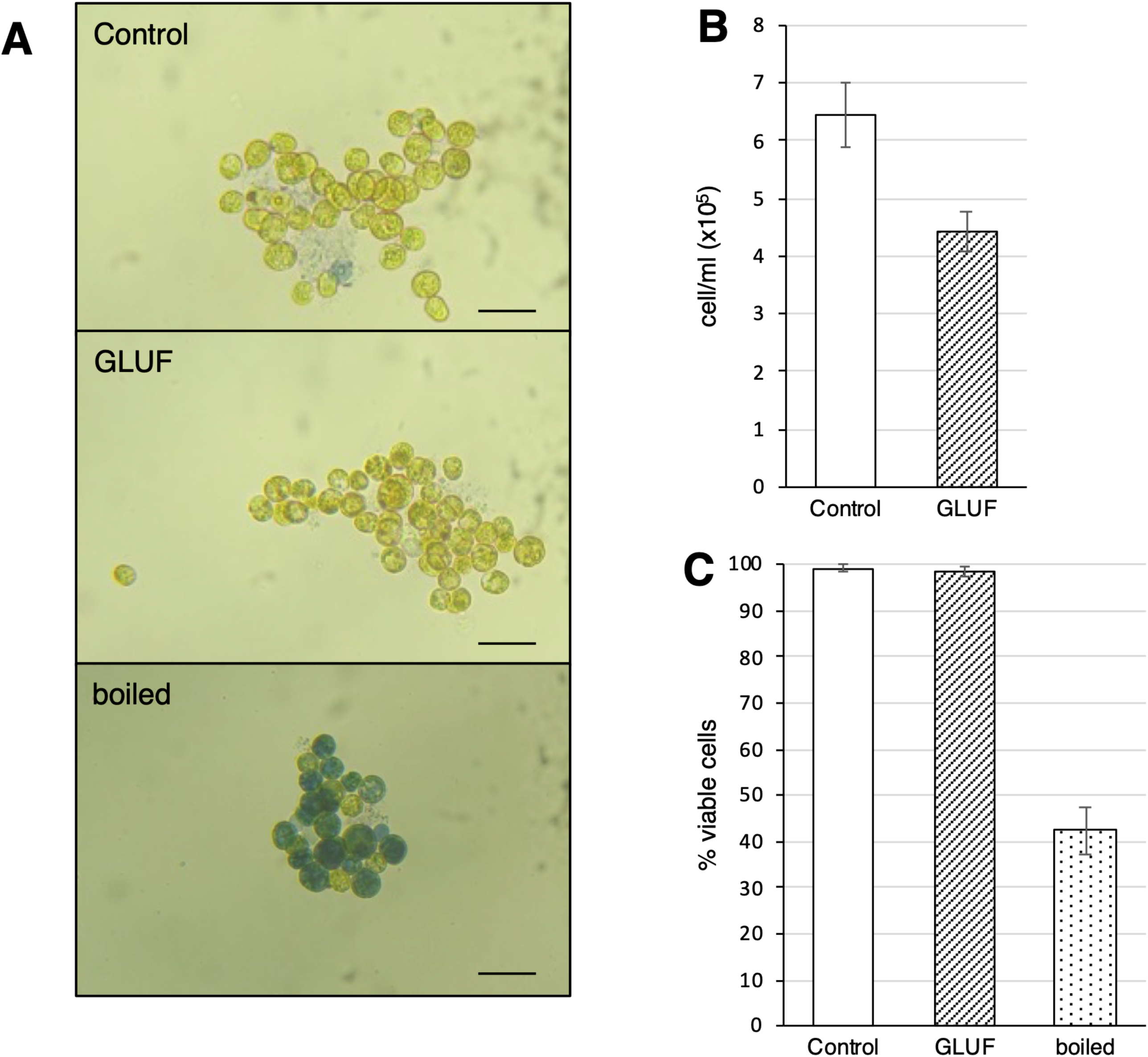
Cell viability in the presence of glufosinate. A. CassKB8 cells after 12 h incubation under darkness with vehicle (Control) or 1 mg/ml glufosinate (GLUF), stained with Evans blue [13] and showing marginal staining. In contrast, heat-killed cells (boiled) show heavy staining. B. Cell count of the cells after the 12 h incubation period with vehicle (white bar, Control) or glufosinate (hatched bar, GLUF). C. Percentage of viable cells from three replicates of representative samples from cells incubated with vehicle (white bar, Control) or glufosinate (hatched bar, GLUF), or heat-killed (dotted bar, boiled). Maximum values from graphs are ± standard error fo the mean.

The effect of GLUF on SBiP1 phosphorylation was then assessed just prior to the onset of the light phase (Dark 12 h), as well as 30 and 120 min of light. The cells under darkness both treated with vehicle (Control) or GLUF displayed the typical behavior of highly Thr phosphorylated SBiP1 (Fig. 4A, Dark 12 h, left upper panels labeled SBiP1-p, respectively; and Fig. 4B, left black bar, respectively). In the control group, upon light exposure for 30 and 120 min, the expected prompt SBiP1 dephosphorylation occurred, (Fig. 4A, left upper panel (SBiP1-p), 30 and 120 min, respectively; Fig. 4B, white bars, 30 and 120 min, respectively). Conversely, in the presence of GLUF, SBiP1 was not significantly dephosphorylated after 30 min of light (Fig. 4A, right upper panel (SBiP1-p), 30 min; Fig. 4B, diagonally hatched bar, GLUF, 30 min). The SBiP1 phosphorylation signal diminished significantly only after 2 h of light (Fig. 4A, right upper panel (SBiP1-p), 120 min; Fig. 4B, diagonally hatched bar, GLUF, 120 min. These data suggested that inhibition of processes involved in nitrogen assimilation profoundly affected the SBiP1 chaperone activation status.

**Figure 4.**
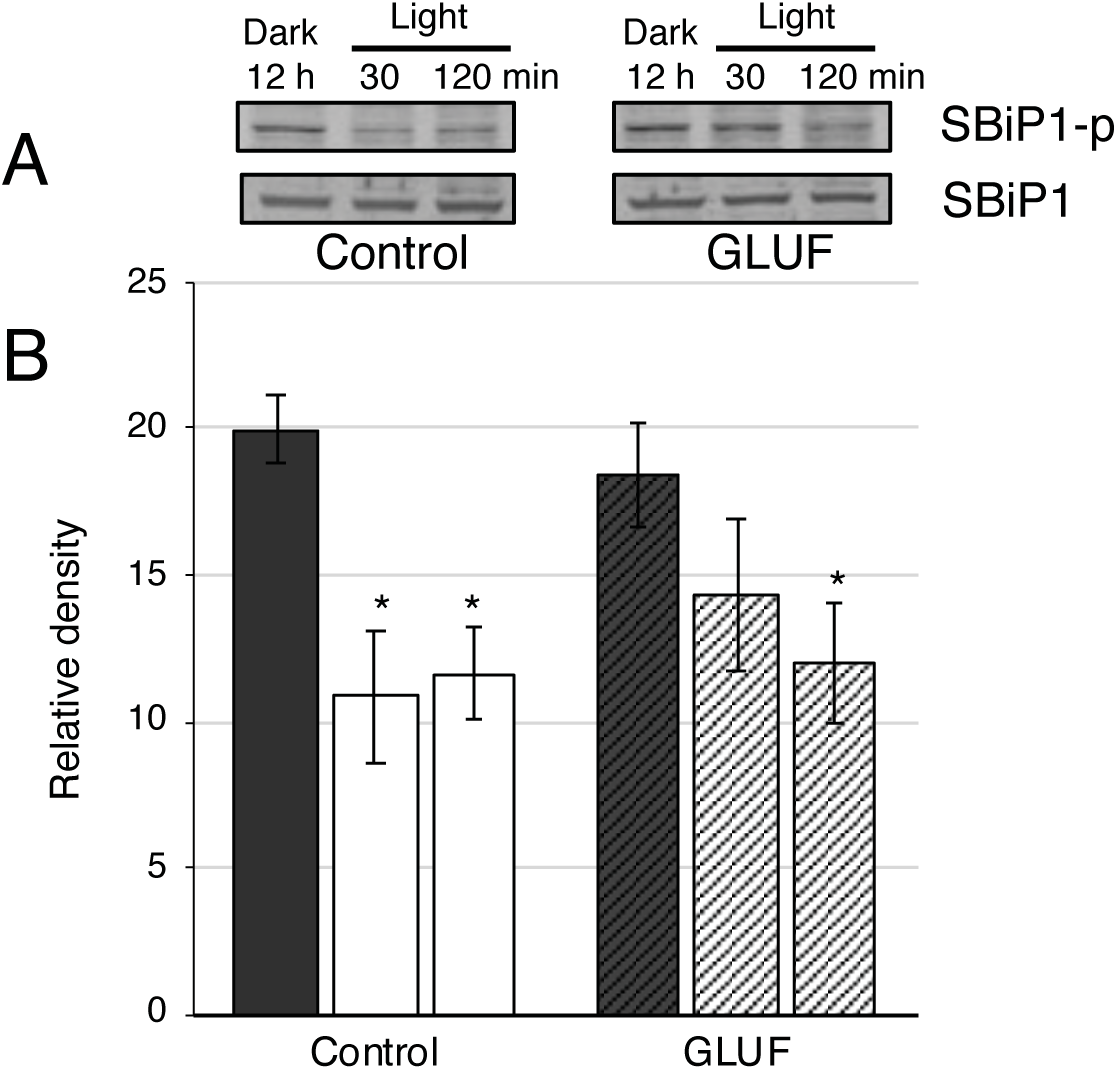
Effect of glufosinate on the Thr phosphorylation behavior of SBiP1 during the dark-to-light transition. A. Upper panels: representative western blots showing the level of SBiP1 Thr phosphorylation (SBiP1-p) from dark-adapted CassKB8 cells exposed to 1 mg/ml glufosinate (GLUF), compared with vehicle only (Control) after 30 min, and 120 min of light stimulation. SBiP1 is highly Thr-phosphorylated (SBiP1-p) after 12 h of continuous darkness in the presence of either vehicle (SBiP1-p, left panel, Dark, 12 h) or 1 mg/ml GLUF (SBiP1-p, right panel, Dark, 12 h). SBiP1 phosphorylation levels diminish after 30 and 120 min of light stimulation in the presence of vehicle (SBiP1-p, left panel, Light, 30 and 120 min, respectively); however, significant SBiP1 Thr phosphorylation is still observed after 30 min of light stimulation in the presence of GLUF (SBiP1-p, right panel, Light, 30 min), but its levels diminish after 120 min (SBiP1-p, right panel, Light, 120 min). Lower panels: western blots of the same treatments and time points analyzed with anti-SBiP1 antibodies (SBiP1) as internal protein loading controls and used for normalization of the densitometric analysis, showing constant levels of the SBiP1 protein throughout the treatments. B. Densitometric analysis of four biological replicates (± standar error of the mean) quantitatively showing the levels of SBiP1 Thr phosphorylation under darkness (black bars) in the absence (Control) or presence of 1 mg/ml GLUF (GLUF), and after 30 min and 120 min of light stimulation in the presence of vehicle (white bars), or GLUF (diagonally hatched bars). Asterisks show statistically significant values compared to the dark condition for each treatment (p<0.05).

## Discussion

HSP70 family chaperones have been reported to be closely linked to cellular metabolism [10,14]. As previously mentioned, in *C. reinhardtii*, the specific inhibition of CrTOR by rapamycin promoted phosphorylation (inactivation) of the ER resident BiP chaperone (CrBiP) and assigned a role to TOR on the control of BiP modification [10]. The CrBiP chaperone inactivation effect was also promoted by treatment with CHX, implicating that CrTOR regulates CrBiP phosphorylation through controlling protein biosynthesis [10]. These data suggested that a close link between the chaperone activation switch and the metabolic status of the cell exists, although the external stimulus that triggers the process remains unknown. We previously found that light profoundly affects Thr phosphorylation of SBiP1 in CassKB8 cells [4–6], and here we demonstrate that a protein synthesis-SBiP1 Thr phosphorylation status correlation occurs similar to CrBiP [10]. We observed that under light conditions, the inhibition of protein synthesis with CHX increased the level of SBiP1 phosphorylation (Fig. 1). Additionally, the dark-adapted CassKB8 cells in the presence of CHX, did not show an observable change in the Thr phosphorylation status of SBiP1 after 60 min of light stimulation (Fig. 2, CHX), contrary to the untreated control (Fig. 2, EtOH). These results indicate that inhibition of protein synthesis maintains the phosphorylated SBiP1 status which renders the chaperone inactive and suggests that protein biosynthesis is the process through which light induces dephosphorylation of SBiP1 which represents its active state. At present, it is unknown whether the TOR pathway is involved in regulation of the chaperone activity as reported for *C*. *reinhardtii* [10]. Upon BLAST analysis to search for TOR in Symbiodiniaceae databases, we could not find any significant homologous sequence, suggesting that a canonical TOR homolog is not present in these organisms and alternate pathways are at play. On the other hand, as expected for a chaperone, the SBiP1 intracellular inactive status by CHX was reversed upon heat shock; that is, the CHX treated cells after 60 min of light stimulation further treated for 30 min with high temperature prompted the activation of the chaperone, observed as a significant dephosphorylation (Fig. 2, CHX, Heat Shock). A previous similar result was found when highly phosphorylated (inactive) SBiP1 under dark conditions was activated by dephosphorylation with the same heat shock treatment [5]. These results indicate that beyond the activation of SBiP1 by light through protein synthesis, heat shock activates SBiP1 by an unknown alternate mechanism, possibly to assist heat-induced unfolded proteins at the ER. It is important to note that inhibition of cytosolic protein synthesis by CHX was concomitant to inactivation of the chaperone whereas inhibition of chloroplastic protein synthesis by CPL had no effect as activation (i.e., Thr dephosphorylation) of SBiP1 occurred analogous to control (Fig. 1). These data suggest that global protein synthesis reflecting the intracellular nutritional status, is closely linked to the chaperone activation switch. Protein synthesis is an anabolic process and similar converging pathways are likely to have impact on regulation of the chaperone activity. Thus, inhibition of another metabolic pathway should result in a similar inactivation of the chaperone. Glutamine synthetase (GS) is a fundamental enzyme for nitrogen assimilation and glutamine synthesis in photosynthetic organisms [15,16]. In Symbiodiniaceae, as well as in plants, inhibition of GS by GLUF inhibits growth (Fig. 3) and the cells eventually perish [17,18]; however, cell death does not occur immediately, and cells are still viable during the first 12 h of GLUF treatment (Fig. 3). Therefore, we could still observe short-term effects if any of, inhibition of GS on SBiP1. Interestingly and consistent with the effect of CHX on the cells, GLUF treatment for 12 h in dark-adapted cells subsequently stimulated with light for 30 min, also appeared to prevent the dephosphorylation of SBiP1 in order to keep the chaperone inactive (Fig. 4A and B, GLUF, 30 min). The effect of GLUF inhibition of SBiP1 Thr dephosphorylation appeared to be transient, as dephosphorylation was observed after 2 h of light stimulation (Fig. 4, GLUF, 120 min), compared to the untreated control (Fig. 4A and B, Control, 120 min).

Given that: a) the GS/GOGAT pathway is the most expedite for ammonium assimilation into aminoacids, and this is the main pathway in dinoflagellates [19,20]; b) light improves ammonium and soluble aminoacid assimilation [21,22,23]; and c) SBiP1 participates in *de novo* protein synthesis, altogether suggests a scenario where a GLUF-induced delay on SBiP1 activation occurs. This is likely due to a downshift in ammonium assimilation which translates into a temporal deficit in protein synthesis until an alternative nitrogen arrival happens through a catabolic pathway such as the pathway through glutamine dehydrogenase which also exists in Symbiodiniaceae [24]. Thus, inhibition of GS had a clear effect on the activation of the chaperone by light. Altogether, the data further supported the findings that inhibition of metabolic processes such as protein and glutamine synthesis have an observable effect on the SBiP1 chaperone activation switch. These results are also consistent with the notion that chaperones of the HSP70 family are influenced by the nutritional status of the cell and this seems to be the case also in CassKB8 cells and probably in Symbiodiniaceae since SBiP1 appears to be a ubiquitous protein in this family [6].

The findings on cultured Symbiodiniacea cannot be extrapolated to symbionts *in hospite*; however, it has become more evident that nitrogen metabolism is critical for symbiosis and the endosymbiot itself. For example, it was recently reported that endosymbionts under optimal light conditions for photosynthesis make more carbon available and stimulate nitrogen assimilation in the host. This in turn, limits nitrogen available to algal endosymbionts resulting in low densities within the holobiont [25]. In fact, it has been demonstrated that cnidarian hosts use a carbon-negative feedback loop to control symbiont proliferation [26] where supplementing additional carbohydrates significantly reduced the symbiont density through limiting their nitrogen availability. Thus, it is quite likely that the light-regulated pathways leading to protein synthesis and nitrogen metabolism in the endosymbiont are coupled to diverse signaling cascades that control the activity of chaperones such as SBiP1 to maintain symbiont-host homeostasis.

A proposed model incorporating our previous data [5,6] and current findings is shown in Fig. 5. Initially, a kinase has rendered SBiP1 fully Thr phosphorylated (thus inactive; light green croissant) under darkness at the normal 26°C growth temperature (right purple arrow). At the onset of the light phase, phototransduction activates light-stimulated pathways (left upper dashed green arrows); among these, a high sensitivity photoreceptor (PR) with broad light spectrum sensitivity [6] upstream of SBiP1 triggers the activation of a currently unknown pathway (lower left dashed green arrow) that is related to the activation of metabolic processes such as protein synthesis (right lower black arrow); the signaling cascade eventually results in the activation of a phosphatase (middle dashed and solid green arrows) leading to the dephosphorylation of Thr phosphorylated SBiP1 (light green croissant) to activate its chaperone activity (green croissant). The active chaperone is now able to assist client proteins being newly synthesized in the ER (SBiP1-protein X-ribosome complex). The PR highly sensitive to the broad visible light spectrum would be a requirement for photoreception and phototransduction in cells such as Symbiodiniaceae living in the water column or in symbiosis deep within the tissue of other underwater hosts. This dephosphorylation occurs independent of photosynthesis since CPL (this work) or DCMU [6] do not affect this SBiP1 activation process (upper left red baton and upper dashed green arrow). This also suggests that alternative nutrient sensing pathways independent of photosynthesis including an orthologous TOR pathway may come into play (left thick green arrow); thus, the chaperone activity switch would also be connected to cues reflecting the nutritional status of the cell (i.e., active protein and glutamine biosynthesis) which would result in activation of the chaperone (middle green dashed and solid arrows). On the other hand, inhibition of global protein synthesis by cycloheximide (CHX) (middle horizontal red baton at the left) results in a decrease in the protein biosynthetic process (right upper black arrow) which somehow promotes (or is the result of) chaperone inactivation (upper middle purple arrow). Alternatively, inhibition of glutamine biosynthesis delays but does not block the activation of the pathways that lead to the activation of the phosphatase (left dashed pink batons) which eventually ensue (left pink arrow following the pink batons and middle dashed and solid green arrows) stimulating the phosphatase activity that dephosphorylates SBiP1 (green croissant). It is currently unknown whether protein synthesis is up- or downstream the chaperone activation switch. Stress responses also affect the chaperone status; under heat stress (32°C) either under light (lower vertical solid green arrow) or darkness (right long solid green arrow), the chaperone activity is required prompting the phosphatase to dephosphorylate SBiP1 [6]. Heat shock also prompts activation of the chaperone even when protein synthesis is inhibited (horizontal left red baton and vertical solid green arrow). The UPR (Unfolded Protein Response) and other stress responses resulting in ROS (Reactive Oxygen Species) production trigger mechanisms critical for cell survival and, in some cases, they could also override the light stimulated activation of the chaperone through various alternative mechanisms including inhibition of protein synthesis (lower vertical red baton). We have shed light on possible links between the modulation of chaperone activity by nutritional cues and phototransduction processes in symbiodiniaceae and are currently searching the molecular participants of the pathways. This will lead us to eventually dissect all the components of this light stimulated signaling mechanism in these photosynthetic potentially endosymbiotic microorganisms.

**Figure 5.**
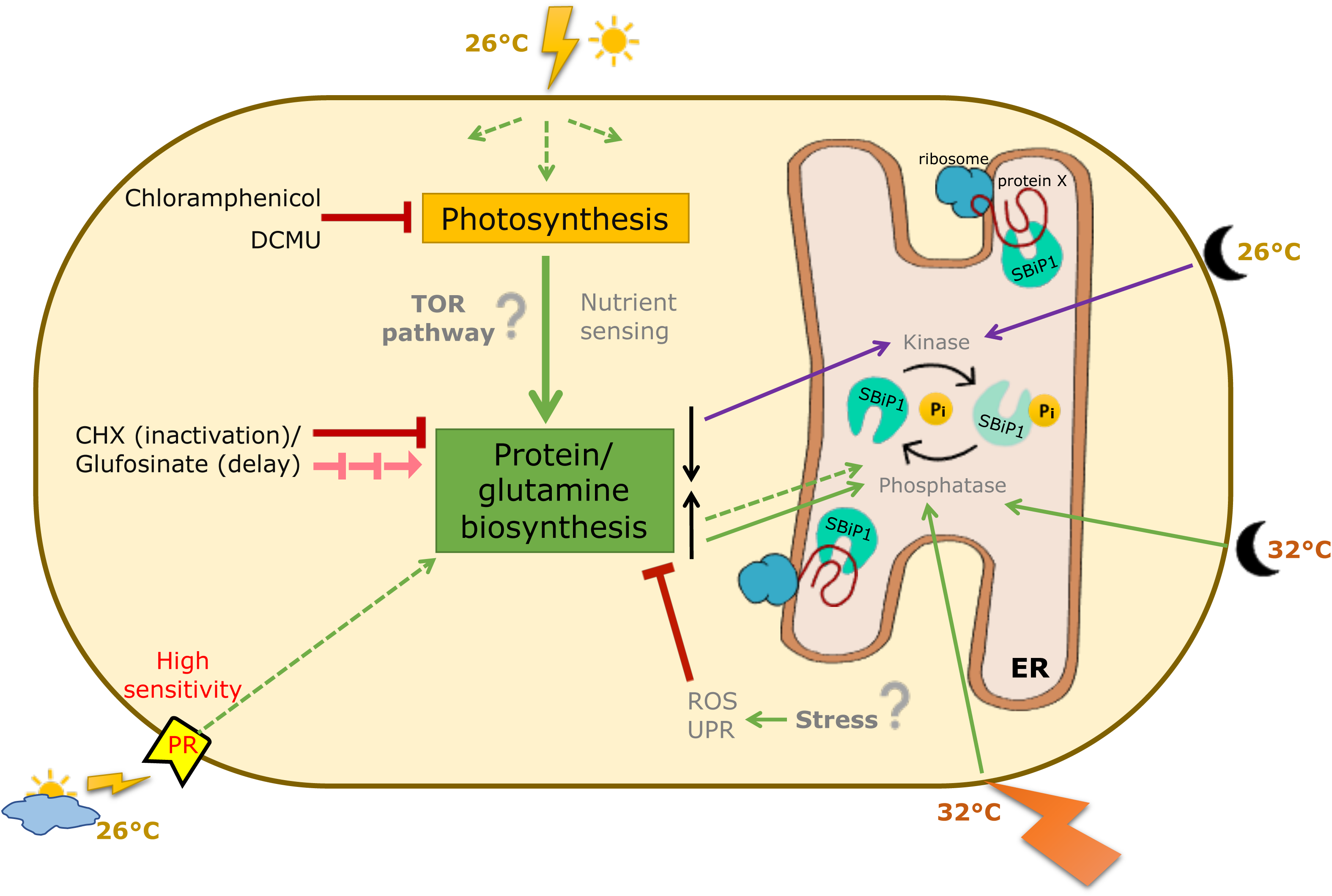
Proposed model of factors that influence the activation/inactivation of SBiP1 chaperone by light. During the dark photoperiod phase, metabolic processes are shut down and SBiP1 is fully phosphorylated (inactive; light green croissant) by an unknown kinase (right upper purple arrow). Upon light exposure, the anabolic pathways activated by phototransduction (upper left dashed green arrows) initiate including photosynthesis (lower left thick green arrow). This causes protein biosynthesis to initiate (thick green arrow at the left and right lower black arrow) and prompts unknown signaling pathways (middle green dashed and solid arrows) that activate the phosphatase causing the chaperone to dephosphorylate and activate (green croissant) to aid in the process. Inhibition of global protein synthesis by cycloheximide (CHX) (middle horizontal red baton at the left) results in a decrease in the protein biosynthetic process (right upper black arrow) which somehow promotes (or is the result of) chaperone inactivation (middle upper purple arrow). Alternatively, inhibition of glutamine biosynthesis delays but does not block the activation of the pathways that lead to the activation of the phosphatase (left dashed pink batons) which eventually ensue (left pink arrow following the pink batons and middle dashed and solid green arrows) and dephosphorylate SBiP1 (green croissant). The effect of light stimulation on SBiP1 inactivation is independent of photosynthesis since it still occurs regardless of its inhibition of PSII by DCMU**^6^** or chloroplast protein synthesis by chloramphenicol (upper red baton). Stress caused by heat shock (32°C) either under light or darkness likely produces ROS and triggers survival mechanisms including the unfolded protein response (UPR) which, in turn, causes activation of the phosphatase (middle lower green arrow) that dephosphorylates and activates SBiP1 (green croissant). The relationship of SBiP1 with anabolic processes may be related to nutrient sensing mechanisms similar to those mediated by the Target of Rapamycin (TOR) pathway. Finally, lower temperature (26°C) does not seem to affect the chaperone activation switch.

## Methods

### CassKB8 cell cultures

CassKB8 culture (also known as MAC-CassKB8 and previously classified as clade A *Symbiodinium*) was a kind gift of Dr. Mary Alice Coffroth (State University of New York at Buffalo). Cultures have been routinely maintained in our laboratory for over twenty years, grown in ASP-8A medium under photoperiod cycles of 12 h light/dark at 26°C at a light intensity of 80-120 μmole photon m^-2^ sec^-1^.

### Antibodies and reagents

Polyclonal anti-phosphothreonine (anti-pThr) antibodies were from Cell Signaling Technology ™, Inc. (Danvers, MA; cat. 9381S) and Abcam (Cambridge, UK; cat. Ab9337). Anti-PsbA to the C-terminal of the PsbA protein (AS05 084) was from Agrisera (Vannas, Sweden). Rabbit anti-SBiP1 polyclonal antibodies were used as the purified IgG fraction (∼ 1 mg/ml IgG as stock) as previously described [5,6]. Alkaline-phosphatase (AP) conjugated polyclonal anti-rabbit IgG antibodies raised in goat were from Zymed^®^-Life Technologies (Grand Island, NY). Reagents 5-bromo-4-chloro-3-indolyl phosphate (BCIP) and nitro blue tetrazolium (NBT) were from Promega (Madison, WI). The inhibitors of protein synthesis cycloheximide (CXH), and chloramphenicol (CLP) were from Sigma. The inhibitor of glutamine synthetase (GS) ammonium glufosinate (GLUF) was from SantaCruz Biothecnology. All other reagents were from Sigma.

### Protein electrophoresis in SDS-PAGE gels and western blot

The protein extracts (see below) were separated in discontinuous denaturing gels [27], of 10 % polyacrylamide in the separation zone [375 mM Tris-HCl, pH 8.8; 10 % acrylamide/bis-acrylamide (29:1); 0.1 % SDS; 0.1 % ammonium persulphate (APS); 0.106 % N, N, N, N’-tetramethylethylenediamine (TEMED)], and 4 % polyacrylamide in the stacking zone [125 mM Tris-HCl, pH 6.8; 4 % acrylamide/bisacrylamide (29:1); 0.1 % SDS; 0.1 % APS; 0.066 % TEMED], in a Mini-PROTEAN^®^3 System (Bio-Rad, Hercules, CA). After electrophoresis, the proteins were transferred to PVDF membranes in “friendly buffer” [28] (25 mM Tris-HCl, 192 mM glycine, 10 % isopropanol) at a constant current of 300 mA for 1 h. The membranes were blocked in a solution of 3 % bovine serum albumin (BSA) in PBS (2.79 mM NaH_2_PO_4_, 7.197 mM Na_2_HPO_4_, 136.9 mM NaCl, pH 7.5) added with 0.01 % TX-100 (PBS-T) for 1 h at 50°C, with gentle agitation. After blocking, the primary antibodies, anti-pThr (Cell Signaling, 1:2,500), anti-PsbA (1:2,500), or anti-SBiP1 (0.404 µg/ml IgG fraction) in PBS-T, were added to the membranes and incubated overnight with gentle rocking at 25°C. Alternatively, when Abcam anti-pThr antibodies were used, TBS-T (20 mM Tris-Base, 150 mM NaCl, 0.01 % TX-100, pH 7.5) was the buffer for the BSA blocking solution (5 % BSA) and primary antibody dilution (1:500). Blocking for at least 2 h and all other incubations were carried out at 4°C. After overnight incubation, the membranes were washed five times, 5 min each, in PBS-T or TBS-T, and incubated with secondary antibody (alkaline-phosphatase conjugated goat anti-rabbit IgG) at 1:2,500 dilution in PBS-T or TBS-T for 2 h at 25 or 4°C, buffer and temperature depending on the primary antibody used. Subsequently, the membranes were washed again five times, 5 min, and the final wash was followed by a brief rinse with alkaline developing solution (100 mM Tris-HCl, 150 mM NaCl, 1 mM MgCl_2_, pH 9) at 25°C. Finally, the membranes were developed with a commercial solution of 5-bromo-4-chloro-3-indolyl phosphate (BCIP) and nitro blue tetrazolium (NBT) according to the manufacturer (Promega, Madison, WI), in alkaline developing solution. In the case of anti-pThr from Cell Signaling, development was carried out in PBS-T to ensure more astringency and less background. It is important to note that we have previously determined that incubations in PBS-T or TBS-T yielded identical results when anti-pThr antibodies from Cell Signaling were used [4].

### Preparation of total protein extracts from Symbiodiniaceae

After each time point of treatments, the Erlenmeyer flask was shaken briefly to suspend the cells and equal 10 ml aliquots were placed into 15 ml falcon tubes. Then, the cells from each tube were sedimented by centrifugation at 2,600 x *g* for 1 min at 26°C, and immediately suspended in 300 μl Laemmli buffer [27], supplemented with 0.2 mM NaVO_3_, 10 mM NaPPi and a cocktail of protease inhibitors (Complete®; Roche, Basel, Switzerland), mixed with ∼ 250 μl total volume of glass beads (425–600 μm diameter). Subsequently, the cells were lysed with a MINI-BEAD BEATER-1® (Biospec Products) at maximum speed for 3 min. The lysate was heated at 95°C for 5 min centrifuged at 12,000 x *g* for 10 min, and the supernatant used for western blot analysis.

### Thr phosphorylation analysis of SBiP1 from CassKB8 cells in the presence of CHX during the transition of darkness to light

Six d-old CassKB8 cultures were sedimented by centrifugation at 2,600 x *g* for 3 min and suspended in fresh ASP-8A medium to reach a concentration of 4-6 x 10^5^ cells/ ml. Two equal portions of 40 ml in Erlenmeyer flasks wrapped with aluminum foil were incubated for 12 h at 26°C under darkness (nocturnal phase). The cultures were then treated 2 h (after 10 h of darkness) before the change to light with either: 0.1 mM CHX (16 µl of 0.25 M CHX stock) in ethanol (EtOH) or with vehicle (16 µl EtOH) as negative control. Protein extraction was carried out at either 12 h darkness, 30, or 60 min of light for each treatment, as described above. For the CHX treatment, a further short-term heat shock was applied after 60 min of light stimulation (see below). The proteins were analyzed by 10 % SDS-PAGE gels, and western blot with either anti-pThr (Abcam), or anti-SBiP1 antibodies. The bands on the blots were analyzed by densitometry and normalized as above. The data from four independent experiments were averaged, subjected to both ANOVA and Student test analyses, and plotted.

### Thr phosphorylation analysis of SBiP1 from CassKB8 cells with CHX or chloramphenicol under continuous light

Six d-old CassKB8 cultures were sedimented by centrifugation at 2,600 x *g* for 3 min and suspended in fresh ASP-8A medium to reach a concentration of 4-6 x 10^5^ cells/ml. Then, three 40 ml portions of cell suspension were separated in Erlenmeyer flasks wrapped with aluminum foil and incubated at 26°C for the 12 h of their nocturnal phase. After this phase, light ensued (80-120 µmol photon m^-2^ s^-1^ and 26°C) and after 3 h, each of the flasks were treated with either 0.1 mM CPL (16 µl of 0.25 M CPL stock in EtOH), 0.1 mM CHX (16 µl of 0.25 M CHX stock in EtOH), or vehicle (16 µl EtOH), as negative control. Prior to protein extraction, the flasks were shaken after each treatment in order to take 10 ml homogeneus aliquots into 15 ml Falcon tubes after 30 min, 1, 4 or 7 h, and the cells were then processed as above. The proteins were analyzed by SDS-PAGE gels, and western blot with either anti-pThr (Cell Signaling), anti-SBiP1, or anti-PsbA antibodies. The bands on the blots were analyzed by densitometry as above. Normalization was carried out with the corresponding immunostained SBiP1 band density. The data from three independent experiments were averaged, subjected to both ANOVA and Student test analyses, and plotted.

### Thr phosphorylation analysis of SBiP1 from dark-adapted cells in the presence of CHX after 60 min light followed by a short-term heat shock

A parallel sample (10 ml suspension) of CassKB8 cells from the CHX treatment (CHX applied 2 h before the dark to light transition), was taken after 60 min of light stimulation at 26°C, centrifuged at 2,600 x *g* for 1 min, resuspended in 10 ml ASP-8A medium with 1 mM CHX and incubated for additional 30 min at 32°C under the same light intensity. After this time, the sample was processed for protein extraction and analyzed as above by SDS-PAGE gels, western blot with either anti-pThr (Abcam), or anti-SBiP1 antibodies, and densitometry as described above.

### Thr Phosphorylation analysis of SBiP1 from cells with glufosinate during the transition of darkness to light

A 6-d-old culture of CassKB8, was concentrated by centrifugation at 2,600 x *g* for 3 min and suspended in 60 ml of ASP-8A medium to reach a cell concentration of 4-6 x 10^5^ cells/ml. The suspension was then divided in 30 ml portions in two separate Erlenmeyer flasks. To one flask, GLUF (ChemCruz®, Santa Cruz Biotechnology, Santa Cruz, CA, USA) was added to a final concentration of 1 mg/ml, and the equivalent amount of vehicle (sterile milliQ water) was added to the other one as control. The flasks were covered with aluminum foil and allowed to incubate for 12 h of their dark photoperiod. In parallel, protein extractions were performed from 10 ml homogeneous aliquots, prior to treatment with light (12 h dark), or after 30- and 120-min light exposure. The samples were analyzed with anti-pThr (Abcam) and anti-SBiP1 antibodies. The assay was performed to four biological replicates, and the bands corresponding to phosphorylated SBiP1 were quantified by densitometry and normalized with the bands of total SBiP1 protein. Band intensities of the phosphorylated SBiP1 were normalized to the corresponding total SBiP1 protein band, and the obtained values averaged and subjected to both ANOVA and Student test analyses and plotted. Cell viability in the presence and absence of GLUF was determined by Evan’s blue staining as previously described [13] under a Leica DM500 microscope.

## Acknowledgements

We are grateful for the technical help of M.O. Edgar Escalante-Mancera, M.I. Miguel Ángel Gómez-Reali, Dr. Edén Magaña-Gallegos, M.S. Fernando Negtete-Soto, M.T.I.A. Gustavo Villarreal-Brito, and M.S. Laura Celis-Gutiérrez.

## Author contributions

RC-M designed and performed experiments, wrote parts of the manuscript; TI-F and EM-R performed data analysis, designed figures, and contributed to the manuscript; MAV developed the concept, designed experiments, wrote the manuscript and obtained funding. All authors read, proofread, and approved the final manuscript.

## Declarations

### Ethical approval

No ethical approval was required for the live materials used in this work since the Symbiodiniaceae cell cultures are publicly available.

### Data availability

All data generated or analysed during this study are included in this article [and its supplementary information files].

