## Supplementary Figure 1 for "Inhibition of protein or glutamine biosynthesis affect the activation of the light stimulated SBiP1 chaperone by dephosphorylation in Symbiodiniaceae"

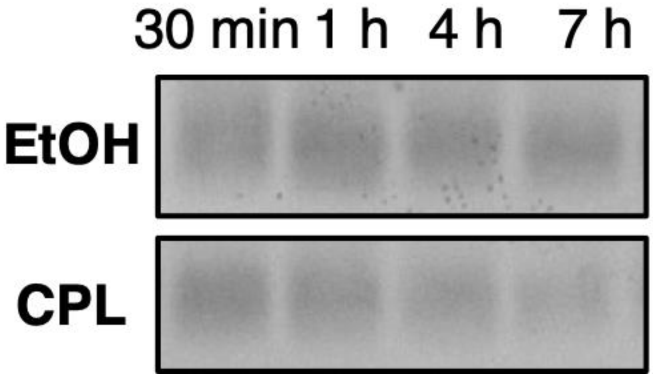


**Supplementary Figure 1.** Effect of chloramphenicol on D1 protein synthesis. Western blot analysis of D1 protein from CassKB8 cells incubated with 0.1 mM chloramphenicol (CPL) or vehicle (EtOH) for 30 min, 1, 4 nd 7 h. The band intensity of D1 clearly diminished after the CPL treatment (CPL; lanes 30 min, 1-7 h), whereas it remained constant along the same time points when vehicle alone was used for the treatment (EtOH; lanes 30 min, 1-7 h).
